# Chronic hyperglycaemia drives functional impairment of lymphocytes in diabetic *INS*^C94Y^ transgenic pigs

**DOI:** 10.1101/2020.08.26.267914

**Authors:** Isabella-Maria Giese, Simone Renner, Eckhard Wolf, Stefanie M. Hauck, Cornelia A. Deeg

## Abstract

People with diabetes mellitus have an increased risk for infections, however, there is still a critical gap in precise knowledge about altered immune mechanisms in this disease. Since diabetic *INS*^C94Y^ transgenic pigs exhibit elevated blood glucose and a stable diabetic phenotype soon after birth, they provide a favourable model to explore functional alterations of immune cells in an early stage of diabetes mellitus *in vivo*. Hence, we investigated peripheral blood mononuclear cells (PBMC) of these diabetic pigs compared to non-transgenic wild-type littermates. We found a 5-fold decreased proliferative response of T cells in *INS*^C94Y^ tg pigs to polyclonal T cell mitogen phytohaemagglutinin (PHA). Using label-free LC-MS/MS, a total of 2,704 proteins were quantified, and distinct changes in protein abundances in CD4^+^ T cells of early-stage diabetic pigs were detectable. Additionally, we found significant increases in mitochondrial oxygen consumption rate (OCR) and higher basal glycolytic activity in PBMC of diabetic *INS*^C94Y^ tg pigs, indicating an altered metabolic immune cell phenotype in diabetics. Thus, our study provides new insights into molecular mechanisms of dysregulated immune cells triggered by permanent hyperglycaemia.

## Introduction

Diabetes mellitus is a major risk factor regarding the outcome of bacterial and viral infections (Harding, Pavkov, Magliano, Shaw, & Gregg, 2019). Among diabetic patients, increased numbers of highly susceptible individuals suffering from tuberculosis (Kumar et al., 2019) or influenza (Marshall et al., 2020) are common. Recently, meta-analyses revealed an enhanced severity and mortality of COVID-19 infections in people with diabetes mellitus (Y. Hu et al., 2020; Zhou et al., 2020). Undoubtedly, there is a link between the disease and an impaired immune system, but the molecular mechanisms involved in altered immune cell function in diabetic patients are still unknown to date. Dysfunctional T cells may play a pivotal role in immunological impairment in diabetes mellitus (Nicholas et al., 2019). However, the precise effects of chronic hyperglycaemia on immune cell function are still uncertain since contradictory findings of both hyperresponsive (Nyambuya, Dludla, Mxinwa, & Nkambule, 2020) and attenuated (Lau et al., 2019; Lee et al., 2019) T cells were reported in type 2 diabetic patients. While numerous studies linked hyperglycaemia to enhanced T cell activation and proliferation (Nyambuya et al., 2020), others suggest that an abnormal glucose homeostasis promotes an insufficient T-cell response via an increased frequency of senescent T cells with proliferative impairment (Lau et al., 2019; Lee et al., 2019). Thus, it has yet to be determined whether dysfunctional T cells are impaired or highly activated in diabetes mellitus and whether hyperglycaemia facilitates immune cell dysregulation. Alterations of cellular mechanisms and molecular networks in disease states can be analysed by changes in proteome composition (Soongsathitanon, Umsa-Ard, & Thongboonkerd, 2019). Still little is known about accurate protein profiles of T cells affected by hyperglycaemia *in vivo*. Therefore, mass spectrometry-based studies are a powerful tool for obtaining comprehensive information regarding cellular pathology (Lepper et al., 2018) and enable hypothesis-generating approaches on the protein level to understand the molecular basis of altered proteome composition and regulation caused by disease. Moreover, recent advances in the field of immuno-metabolism recognized cellular metabolism as a primary driver and regulator of immune cell function, leading to a wide variety of functionally different immune cells (Olenchock, Rathmell, & Vander Heiden, 2017). Since there is still a lack of knowledge about hyperglycaemia-associated alterations of immune cell metabolic phenotypes, there is an urgent need to understand whether and how high glucose levels affect the metabolic phenotype of the cells and thereby change their immune cell function (Olenchock et al., 2017). Diabetic *INS*^C94Y^ transgenic (tg) pigs were generated as a large animal model of permanent neonatal diabetes mellitus (Renner et al., 2013). These pigs are characterized by impaired insulin secretion with consecutive hypoinsulinaemia and increased fasting blood glucose levels (Renner et al., 2013). Early on, *INS*^C94Y^ tg pigs show a stable diabetic phenotype and mirror several disease-associated alterations of diabetes mellitus as seen in humans such as cataract, retinopathy, impaired myocardial function and regeneration (Hinkel et al., 2017; Kleinwort et al., 2017). Since early stages of disease in diabetic patients are often unperceived for a long time, initial immunological alterations are difficult to observe due to the latent disease process. Thus, this diabetic pig model enables the exploration of deviant immune cell function and underlying mechanisms in an early stage of diabetes mellitus with hyperglycaemia and hypoinsulinaemia, offering valuable insights into disease pathogenesis. Hence, we used PBMC of young *INS*^C94Y^ tg pigs and non-transgenic wild-type littermates to investigate the impact of a permanent early-life hyperglycaemic condition on immune cell function.

## Materials and methods

### Animal model and sample preparation

In this study, PBMC of 23 sex-matched diabetic *INS*^C94Y^ tg pigs and 31 non-transgenic wild-type littermates at the age of 12 weeks were used. *INS*^C94Y^ tg pigs were generated previously as described and exhibited significantly elevated random blood glucose levels within 24 h after birth compared with their non-transgenic wild-type littermates (Renner et al., 2013). All animals were housed under controlled conditions, had free access to water and were fed a commercial diet. For sampling, pigs were fasted overnight and heparinized venous whole blood was collected. PBMC were isolated by density gradient centrifugation (RT, 500 x g, 25 min, brake off) with Pancoll separating solution (PAN-Biotech, Aidenbach, Germany), and restored in PBS (pH 7.4) or RPMI medium (PAN-Biotech, Aidenbach, Germany), supplemented with 10% heat-inactivated foetal calf serum (FCS) and 1% penicillin/streptomycin (both Biochrom, Berlin, Germany). Blood withdrawal was performed according to the German Animal Welfare Act with permission from the responsible authority (Government of Upper Bavaria), following the ARRIVE guidelines and Directive 2010/63/EU.

### Lymphocyte stimulation by mitogens

In addition to unstimulated controls, PBMC were either stimulated by pokeweed mitogen (PWM; 1μg/ml), concanavalin A (ConA; 1μg/ml) or phytohaemagglutinin (PHA; 1μg/ml). After an incubation period of 32 hours at 37°C, cells were incorporated with ^3^H-thymidine (Perkin Elmer, Hamburg, Germany) and incubated for 16 more hours. After harvesting, ^3^H-thymidine incorporation was quantified by detecting counts per minute (cpm), using a Microbeta (Perkin Elmer, Hamburg, Germany). Proliferation rate was expressed as factor of ^3^H-thymidine incorporation with respect to unstimulated cells.

### Flow cytometry

Staining of 2×10^5^ cells per well was performed with mouse anti-human CD79α (clone HM57, Bio-Rad AbD Serotec, Puchheim, Germany, 1:100) for identification of B cells and Alexa Fluor 647-conjugated rat anti-human CD3∊ (clone CD3-12, 1:200; cross-reactive to pig (Czajka et al., 2015)) for identification of T cells. We used FITC-conjugated mouse anti-pig CD4 alpha (clone MIL17, Bio-Rad AbD Serotec, Puchheim, Germany, 1:20) and Alexa Fluor 647-conjugated mouse anti-pig CD8α (clone 76-2-11, Becton Dickinson, Heidelberg, Germany, 1:400) to identify αβT cells and mouse anti-pig SWC5 (clone b37c10, Bio-Rad AbD Serotec, Puchheim, Germany, 1:100) for identification of γδT cell subpopulation. If necessary, a secondary antibody was used (Alexa Fluor 647-conjugated goat F(ab’)2 anti-mouse IgG (Fc), Dianova, Hamburg, Germany, 1:1000). Analyses were performed with MACSQuant Analyzer 10 and Flowlogic Software (both Miltenyi Biotec, Bergisch Gladbach, Germany).

### Magnetic activated cell sorting (MACS) for CD4^+^ T cells

Briefly, a total of 6 × 10^7^ cells was incubated in staining buffer with mouse anti-pig CD4 alpha (clone MIL17, Bio-Rad AbD Serotec, Puchheim, Germany, 1:50) at 4°C for 20 min. Staining buffer contained phosphate-buffered saline (pH 7.2) and was supplemented with 2 mM EDTA and 0.5% bovine serum albumin (BSA). In the next step, cells were resuspended in 480 μL staining buffer before adding 120 μL anti-mouse IgG_2a/b_ MicroBeads (Miltenyi Biotec, Bergisch Gladbach, Germany) for an incubation period of 15 min. In further steps, BSA in staining buffer was omitted to prevent interference with mass spectrometry. Magnetic separation was performed using LS columns (Miltenyi Biotec, Bergisch Gladbach, Germany). Magnetically-labelled CD4^+^ T cells were retained in the magnetic field, while unwanted cells were eliminated by three washing steps. Positive CD4^+^ T cell fraction was eluted by removing the column from magnetic field and flushing with staining buffer. 6 × 10^5^ positive selected cells were pelleted and stored at −20 °C until filter-aided sample preparation (FASP). The isolation of porcine CD4^+^ T cell routinely achieved >90% purity, confirmed by flow cytometry.

### Mass spectrometry and data analysis

Purified CD4^+^ T cells of four diabetic *INS*^C94Y^ tg pigs and five littermate wildtypes were analyzed. CD4^+^ T cell pellets were lysed in urea buffer (8M in 0.1M Tris/HCl pH 8.5) and 10 μg total protein of each sample was proteolysed with LysC and trypsin by a modified filter-aided sample preparation (FASP) as described (Grosche et al., 2016). Acidified eluted peptides were analyzed in the data-independent acquisition mode on a Q Exactive HF-X mass spectrometer (Thermo Fisher Scientific, Waltham, MA, USA) online coupled to an ultra-high-performance liquid chromatography (UHPLC) system (Ultimate 3000, Thermo Fisher Scientific). The spectral library was generated directly in Spectronaut Pulsar × (Biognosys, Schlieren, Switzerland; version 12.0.20491.17.25792) as described (Singh et al., 2020). Spectronaut was equipped with the Ensembl Pig Database (Release 75 (Sscrofa10.2), 25,859 sequences, https://www.ensembl.org). Peptide identification was filtered to satisfy an FDR of 1% by the mProphet approach with q-value cut-off at 0.05 (Reiter et al., 2011). Statistical analysis was performed on log2 transformed normalized abundance values using Student’s *t*-test. All proteins with p < 0.05 were considered significantly different and proteins with *INS*^C94Y^/wt ratio ≥ 2.0 were determined as differentially abundant. The heatmap of hierarchical cluster analysis was created with open source software Cluster 3.0 and was illustrated via Java TreeView (version 1.1.6r4, http://jtreeview.sourceforge.net). GraphPad Prism Software (version 5.04) was used to design Volcano plot, and pathway enrichment analysis was done with open source software Reactome (Pathway Browser version 3.6, https://reactome.org).

### Quantification of ANXA1 by immunofluorescence staining

To quantify ANXA1 in porcine PBMC, flow cytometric analysis was performed using rabbit anti-human ANXA1 (Thermo Fisher Scienific, Ulm, Germany, 1:100) and Alexa Fluor 647-conjugated goat anti-rabbit IgG H+L (Invitrogen, Karlsruhe, Germany; 1:500). Cross reactivity of anti-human ANXA1 was proved and high sequence homology of porcine ANXA1 was confirmed via BLASTP (https://blast.ncbi.nlm.nih.gov/Blast.cgi). For intracellular staining, PBMC were permeabilized (BD Cytofix/Cytoperm, Becton Dickinson, Heidelberg, Germany) for 20 min at 4°C and washed with washing buffer (BD Perm/Wash, Becton Dickinson, Heidelberg, Germany), diluted in PBS 1:10. For immunofluorescence staining, purified CD4^+^ T cells of wild-type and *INS*^C94Y^ tg pigs were incubated with FITC-conjugated anti-pig CD4 alpha before staining of ANXA1. After fixation with 1 % PFA, cell nuclei were counterstained with 4′,6-diamidino-2-phenylindole (DAPI; Invitrogen, Karlsruhe, Germany, 1:100) for 30 min at RT. 5 × 10^4^ cells were transferred to microscope slides and centrifugated (300 × g, 10 min) before coverslip using mounting medium (Serva, Rosenheim, Germany). Visualization of stained targets was performed using a Leica Dmi8 microscope with associated LAS-X-software (Leica, Wetzlar, Germany).

### Measurement of OCR and ECAR by Seahorse XFe Analyzer

Metabolic phenotypes of porcine PBMC were determined using a Seahorse XFe Analyzer (Agilent Technologies, Waldbronn, Germany) measuring oxygen consumption rate (OCR), which is attributed to mitochondrial respiration and extracellular acidification rate (ECAR), which can be related to glycolysis (van der Windt, Chang, & Pearce, 2016). In accordance with the manufacturer’s instructions, sterile XF assay buffer (Seahorse XF RPMI Medium supplemented with 10mM glucose, 2mM L-glutamine, and 1mM pyruvate, pH 7.4; Agilent Technologies, Waldbronn, Germany) was used for experiments. Prior to the start of the assay, sensor cartridges (Agilent Technologies, Waldbronn, Germany) were prepared adding oligomycin, FCCP and rotenone & antimycin A. A total of 1 × 10^6^ PBMC was seeded in 24-well XF24 cell culture microplates (Agilent Technologies, Waldbronn, Germany), while four wells were kept free from cells as background correction. Baseline OCR and ECAR were measured before adding oligomycin, FCCP and rotenone & antimycin A. OCR was reported in units of pmol/minute and ECAR in mpH/minute.

### Statistical Analysis

Kolmogorow-Smirnov (KS) test was performed for determination of Gaussian distribution. If KS test indicated p < 0.05 (no normal distribution), Mann-Whitney U test was used for statistical analysis, while student’s *t*-test was used if KS test was p > 0.05 (normal distribution). In both tests, statistical probabilities were considered significant at p < 0.05. Significances are indicated by asterisks with *p < 0.05, **p < 0.01 and ***p < 0.001.

## Results

### Reduced proliferative capacity of lymphocytes in diabetic *INS*^C94Y^ transgenic pigs after polyclonal stimulation with PHA

For assessment of the proliferation response *in vitro*, we used B- and T cell mitogen PWM (Nam et al., 2019) and the specific T cell mitogens ConA (Rodríguez-Gómez et al., 2019) and PHA (Lin, Chen, Liao, Lee, & Chien, 2016) to stimulate lymphocytes of diabetic *INS*^C94Y^ tg pigs and wild-type littermates. Interestingly, while no differences could be observed between the two groups after stimulation with PWM (Fig 1a) and ConA (Fig 1b), lymphocytes of diabetic pigs (n=11) revealed a significantly 5-fold decreased proliferation rate in response to PHA compared to wildtypes (n=13) (Fig 1c; ***p≤0.001).

**Fig. 1.**
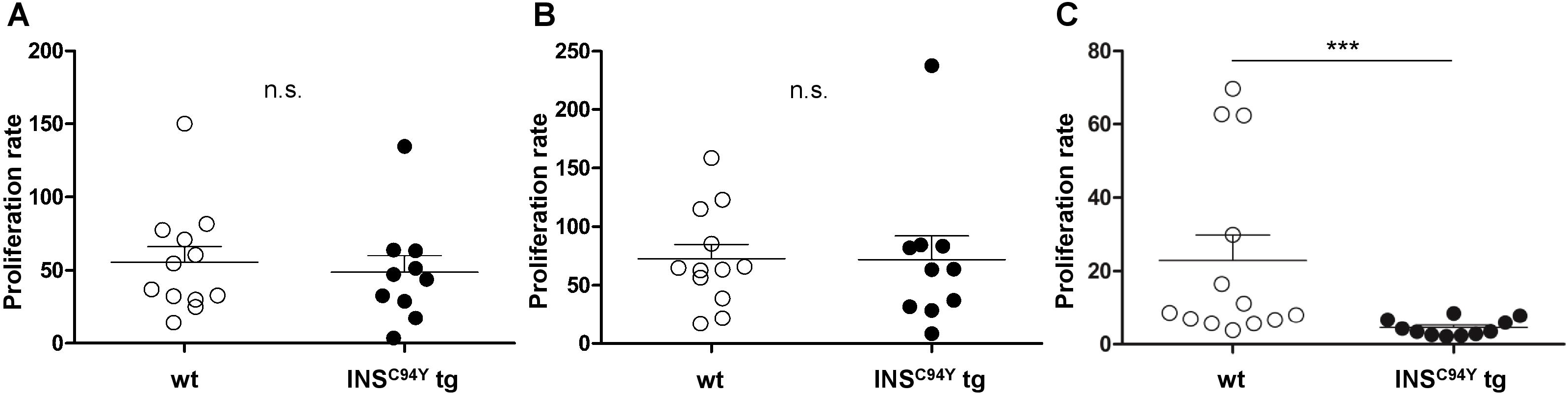
Proliferative capacity of porcine lymphocytes. Mitogens were used for stimulation of lymphocytes analyzed after 48 h of culture: **(A)** pokeweed mitogen (PWM) **(B)** concanavalin A (ConA) or **(C)** phytohaemagglutinin (PHA). Proliferation rate of stimulated lymphocytes did not differ between wild-type (wt, n=12) and *INS*^C94Y^ tg pigs (n=10) stimulated by ConA and PWM. After PHA-stimulation, lymphocytes of *INS*^C94Y^ tg pigs (n=11) revealed a significantly 5-fold decreased proliferation rate (***p≤0.001) compared to non-transgenic wild-type littermates (n=13). Data are represented as means ± SEM; horizontal bars indicate group means.

### Lymphocyte subpopulation ratio did not differ between wildtypes and diabetic pigs

Next, we examined whether the decreased ability to proliferate correlated with altered lymphocyte subpopulations in diabetic *INS*^C94Y^ tg pigs. There were no significant differences in the percentage of B or T cell population between wildtypes and diabetic pigs (Supplementary Fig 1a, b). Moreover, measurements of T cell subsets CD4^+^ (Supplementary Fig 1c), CD8α^+^ (Supplementary Fig 1d), CD4^+^CD8α^+^ αβT cells (Supplementary Fig 1e) and SWC5^+^ γδT cells (Supplementary Fig 1f) revealed no differences between both groups.

### Proteomic profiling identified disease-associated protein ANXA1 in CD4^+^ T cells of diabetic *INS*^C94Y^ transgenic pigs

We characterized the proteomes of porcine CD4^+^ T cells, which we hypothesized to be the key drivers of impaired proliferation response. Purified CD4^+^ T cells were examined with differential proteome analyses, using data-independent acquisition LC−MS/MS. A high-resolution proteome with 2,704 identified proteins was obtained (quantified with at least two unique peptides; Supplementary Table 1), with 80 proteins significantly different in abundance (*p < 0.05; Fig 2a). Two of these proteins had a ≥ 2-fold difference in abundance, named MYL4 and ANXA1 (Fig 2b). In the whole sample set, ANXA1 was highlighted with highest statistical significance in protein abundance differences between wildtypes and diabetic pigs (***p < 0.001). Interestingly, higher levels of ANXA1 were also already described in serum and plasma of type 1 (Purvis et al., 2018) and II diabetes mellitus patients (Gareth S. D. Purvis et al., 2019). Therefore, we subsequently focused on ANXA1 to get further information.

**Fig. 2.**
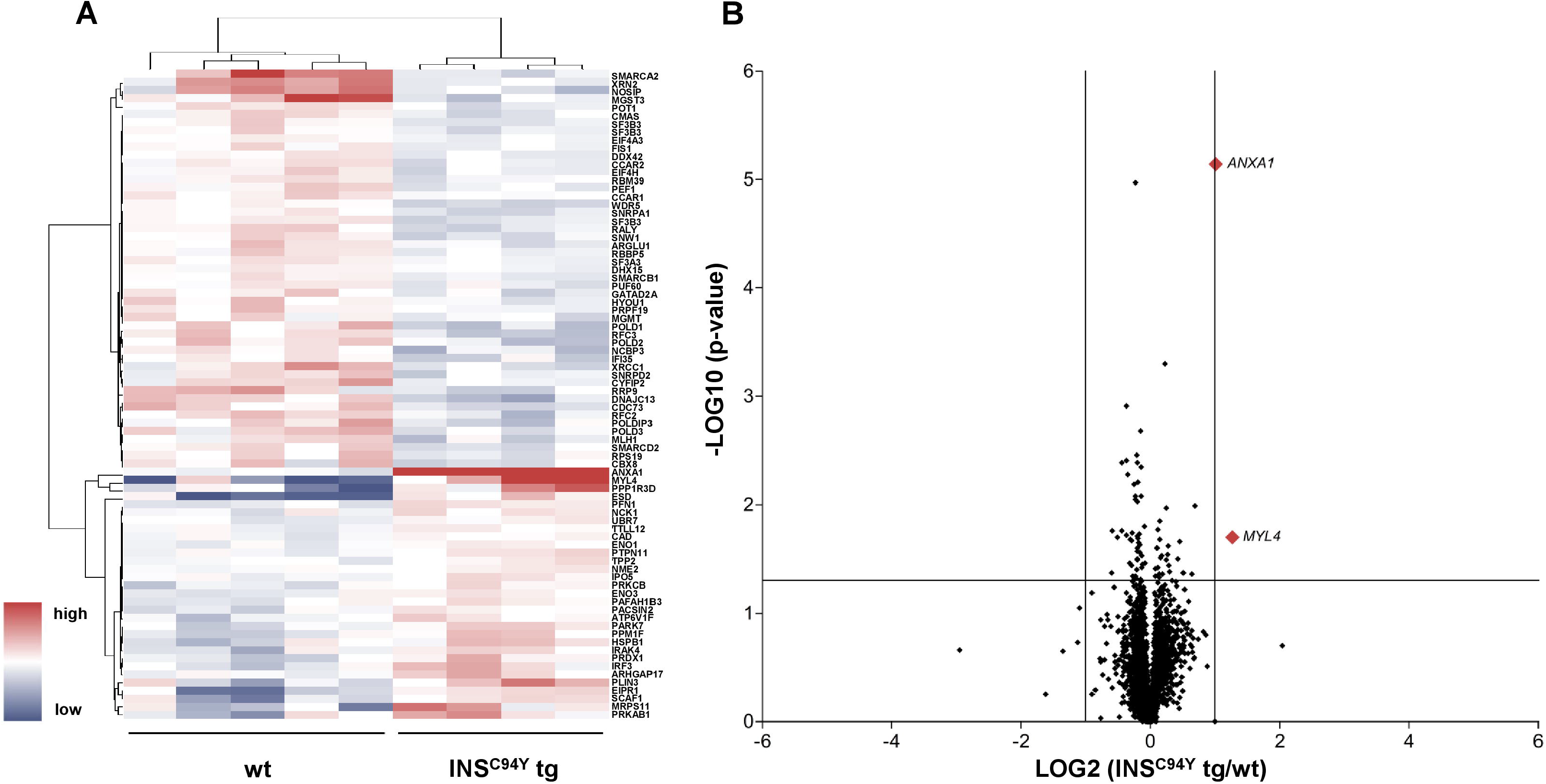
Quantitative proteome analyses of porcine CD4^+^ T cells. **(A)** Hierarchical cluster analysis of the porcine CD4^+^ T cell proteome revealed significant differences in protein abundance (p < 0.05) in *INS*^C94Y^ tg pigs (n=4) and non-transgenic littermates (wt, n=5) indicated by their human orthologue gene names. Red color indicates increased and blue color decreased abundances of the identified proteins. **(B)** Volcano plot illustrates quantified proteins (≥2 unique peptides) in CD4^+^ T cells of *INS*^C94Y^ tg pigs and wild-types (wt) plotted against the respective statistical significance. The horizontal line indicates the p-value cut-off at 0.05. Proteins with a mean fold change ≥ 2 and p <0.05 are shown in red (MYL4 and ANXA1).

### ANXA1 was higher abundant at CD4^+^ T cell outer cell membranes of diabetic *INS*^C94Y^ transgenic pigs

Flow cytometric analyses confirmed that ANXA1 expression was significantly increased in CD4^+^ T cells of diabetic *INS*^C94Y^ tg pigs (n=9) compared to wildtypes (n=11) (*p ≤ 0.05; Fig 3 a-c). Furthermore, we analysed ANXA1 subcellular localization in purified CD4^+^ T cells with immunocytology and flow cytometry. While intracellular ANXA1 levels were equivalent between both groups (Fig 3 d-f; n= 4 per group), ANXA1 was significantly enhanced at the outer cell membrane of CD4^+^ T cells from diabetic pigs (*p<0.05; Fig 3 g-i; n= 6 per group).

**Fig. 3.**
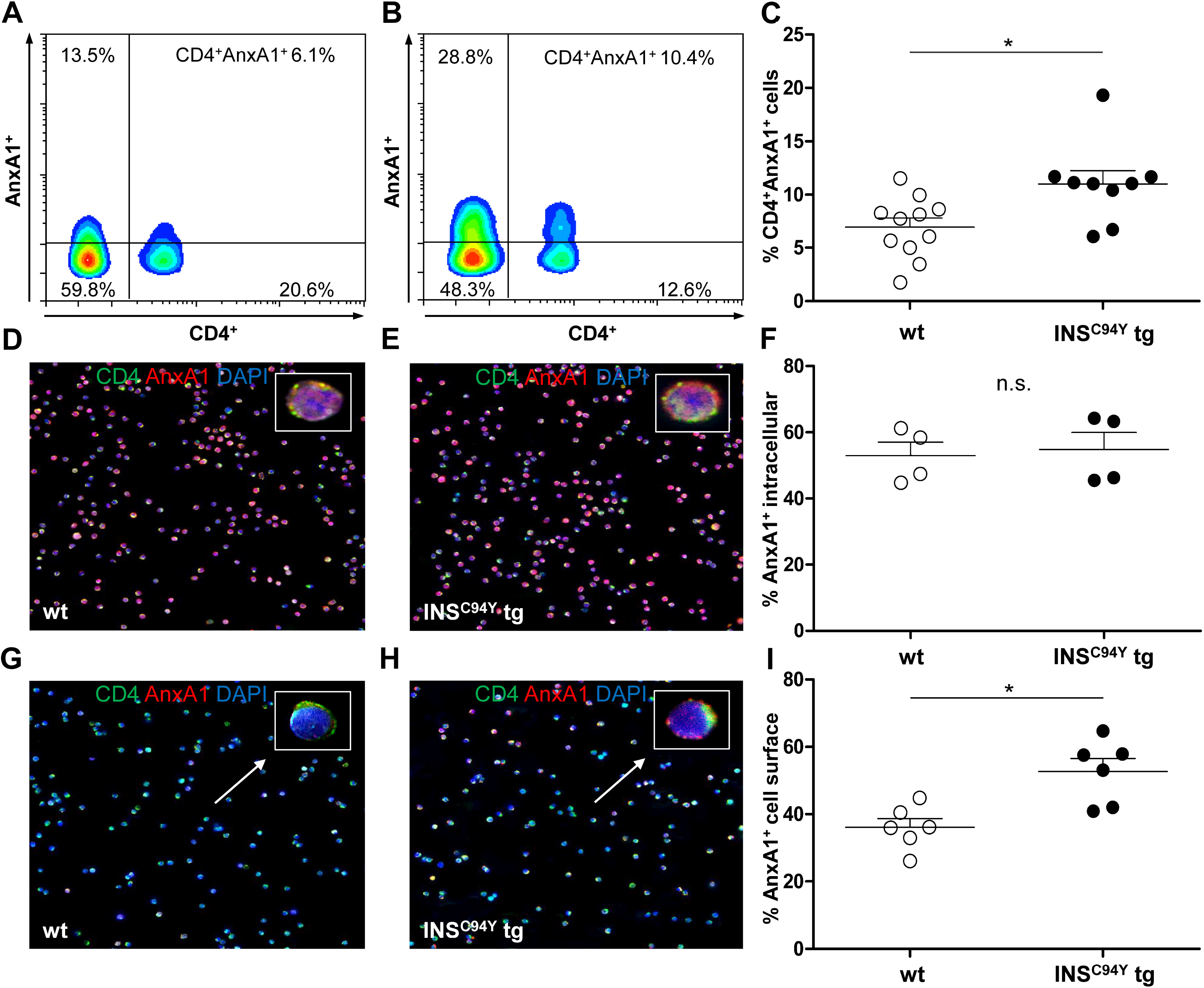
Differential ANXA1 expression in porcine CD4^+^ T cells. **(A-C)** Representative flow cytometric images of ANXA1^+^CD4^+^ T cells of a wild-type **(A)** and an *INS*^C94Y^ tg pig **(B)** are shown with reduced ANXA1 expression in wild-type CD4^+^ T cells. **(C)** Scatter plots summarising flow cytometry analyses of PBMC of eleven wild-type and nine *INS*^C94Y^ tg pigs. ANXA1 was significantly elevated in CD4^+^ T cells of *INS*^C94Y^ tg pigs (*p < 0.05). **(D-F)** Similar intracellular ANXA1 levels in purified CD4^+^ T cells of a wild-type **(D)** and an INS^C94Y^ tg pig **(E)** shown by immunofluorescence staining for ANXA1 (red), CD4 (green) and nucleus (blue, counterstained with DAPI) and confirmed by flow cytometry using n=4 per group **(F)**. **(G-I)** Representative images of immunofluorescence staining for ANXA1 on the outer cell membrane of porcine CD4^+^ T cells demonstrate higher ANXA1 expression in CD4^+^ T cells of the *INS*^C94Y^ tg pig **(H)** compared to the wild-type **(G)** (arrows). **(I)** A higher ANXA1 expression on the cell surface of diabetic CD4^+^ T cells compared to CD4^+^ T cells of wild-types was confirmed by flow cytometry (n=6 per group). Horizontal bars indicate group means.

### Pathway enrichment analyses of CD4^+^ T cells highlight lipophagy pathway in the diabetic condition

To detect altered biological pathways in diabetic immune cells, we performed pathway enrichment analyses via the Reactome database (Jassal et al., 2020). The most significant pathways enriched in CD4^+^ T cells of wildtypes were “Lagging strand synthesis”, “Gap-filling DNA repair synthesis and ligation in GG-NER“ and “Polymerase switching” (Table 1). In contrast, proteins with significantly higher abundances in CD4^+^ T cells of *INS*^C94Y^ tg pigs were associated with the pathways “Downstream signal transduction”, “RUNX1 regulates transcription of genes involved in differentiation of myeloid cells” and “Lipophagy” (Table 2). Since lipophagy is linked to metabolic dysfunction of immune cells (Kounakis, Chaniotakis, Markaki, & Tavernarakis, 2019; Tian et al., 2019), and it was recently shown that functional different cell subtypes can be identified solely by their metabolic phenotypes (Olenchock et al., 2017), we next analysed metabolic profiles of the T cells.

**Table 1.** Pathway Enrichment Analysis of CD4^+^ T cells from wild-type littermates.

| **Most significant pathways enriched in CD4<sup>+</sup> T cells of wild-types** |  |  |
| --- | --- | --- |
| **Pathway name** | **p-value** | **Identified proteins** |
| Lagging Strand Synthesis | 2.00e-12 | POLD1, POLD2, POLD3,<br>RFC2, RFC3 |
| Gap -filling DNA repair<br>synthesis and ligation in<br>GG-NER | 2.69e-12 | XRCC1, POLD1,POLD2,<br>POLD3, RFC2, RFC3 |
| Polymerase switching | 4.45e-12 | POLD1,POLD2, POLD3,<br>RFC2, RFC3 |

**Table 2.** Pathway Enrichment Analysis of CD4^+^ T cells from *INS*^C94Y^ tg pigs.

| Most significant pathways enriched in CD4<sup>+</sup> T cells of *INS<sup>C94Y</sup>* tg pigs |  |  |
| --- | --- | --- |
| Pathway name | p-value | Identified proteins |
| Downstream signal transduction | 4.37e-04 | NCK1, PTPN11 |
| RUNX1 regulates transcription of genes involved in differentiation of myeloid cells | 8.92e-04 | PRKCB |
| Lipophagy | 0.001 | PLIN3, PRKAB1 |

### Increased basal glycolytic activity and mitochondrial respiration in PBMC of diabetic pigs

Interestingly, the basal glycolytic rate of PBMC from diabetic pigs was significantly higher compared to wild-type PBMC (Fig 4a; *p<0.05), indicating increased lactate production and glycolytic activity of diabetic PBMC. Furthermore, analyses of OCR revealed a significant increase in oxygen consumption at all measured time points in diabetic pigs. Compared to wild-type PBMC, a rise of basal, ATP-linked and maximal mitochondrial respiration (**p<0.01) as well as spare respiratory capacity (*p<0.05) was present in PBMC from diabetic pigs (Fig 4b). These differences indicated a fundamentally different metabolic phenotype of PBMC from wildtypes (n=15) and diabetic *INS*^C94Y^ tg pigs (n=10).

**Fig. 4.**
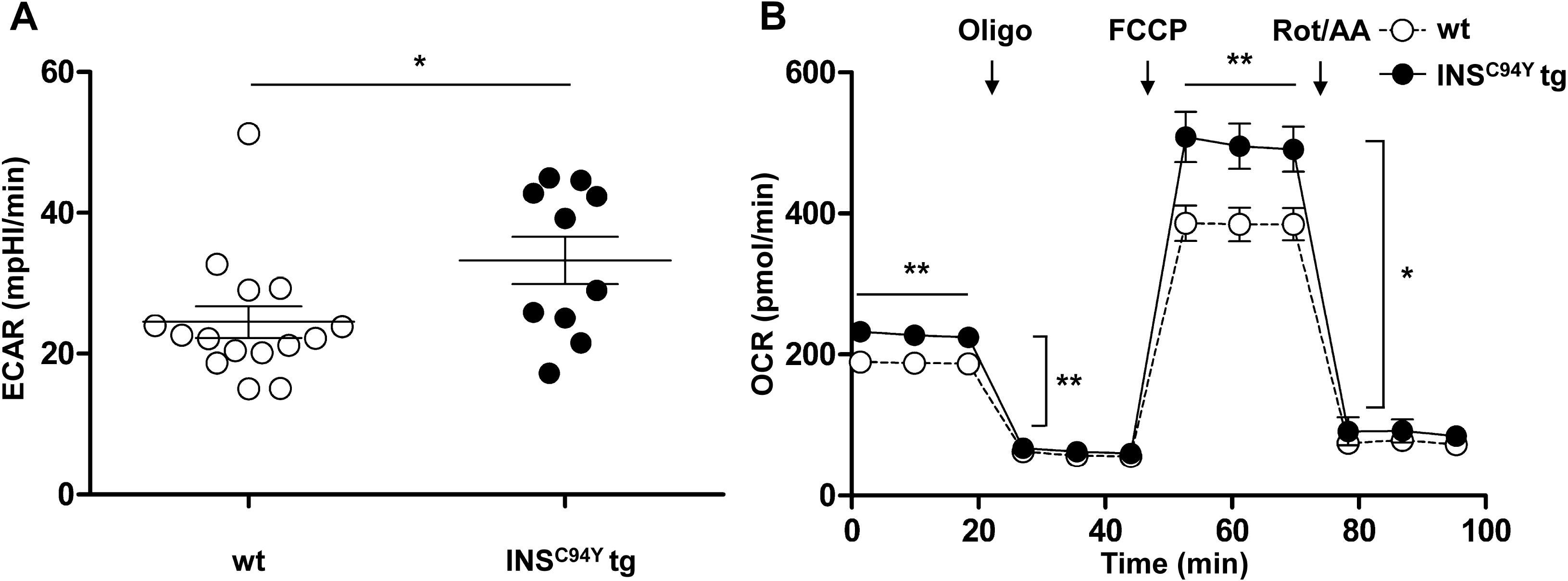
Basal glycolytic and mitochondrial respiratory profiles of porcine PBMC. **(A)** Basal glycolytic rate (Extracellular acidification rate, ECAR) was significantly elevated in PBMC of *INS*^C94Y^ tg pigs (n=10) compared to wild-type littermates (wt, n=15) (*p<0.05, Student’s *t* test) **(B)** Oxygen consumption rate (OCR) was measured under basal conditions and after injection of oligomycin, FCCP, rotenone and antimycin A. Compared to PBMC from wild-types (wt, n=15), mitochondrial respiration of PBMC from diabetic pigs (n=10) yielded a significantly increased basal, ATP-linked and maximal respiration (**p<0.01) as well as a significantly increased spare respiratory capacity (*p<0.05). Data are represented as means ± SEM.

## Discussion

To assess the impact of permanent hyperglycaemia on immune cell function *in vivo*, we characterised lymphocytes of a transgenic pig model for permanent neonatal diabetes mellitus. Significant differences in proliferative response, CD4^+^ T cell proteomes and the metabolic phenotypes were already detectable in these early diabetic pigs. Our findings point to main distinct changes in the T cell compartment triggered by early-life high blood glucose and indicate that several immune alterations already occur in the early stage of diabetes mellitus.

We observed no differences of lymphocyte proliferation between controls and diabetic pigs after polyclonal stimulation with B and T cell mitogen PWM and T cell mitogen ConA. However, after polyclonal stimulation with T cell mitogen PHA, we detected a distinct decreased proliferation of lymphocytes from diabetic pigs, which points to an impaired immune response selectively induced by this mitogen. Interestingly, a decreased response of T cells to PHA-stimulation was already demonstrated in diabetic patients (Chang & Shaio, 1995; Richard et al., 2017). The significantly reduced proliferation to PHA was correlated with a decreased interleukin (IL)-2 production (Richard et al., 2017). The cytokine IL-2, as well as its IL-2 receptor, is a cytokine crucial for proliferation processes in human T cells (Abbas, Trotta, D, Marson, & Bluestone, 2018) and, thus, is enhanced in these cells after PHA stimulation leading to cell proliferation (S. Hu et al., 2019). These findings were supported and extended by another transcriptomic study of PHA-activated human CD4^+^ and CD8^+^ T cells with decreased gene expression of IL-2 and IL-2 receptor in diabetic type 2 patients (Stentz & Kitabchi, 2007). Furthermore, T cells from these patients exhibited differences in expression of additional genes and gene products like lower expression of genes of insulin signalling pathway and enzymes of the glycolytic pathway (Stentz & Kitabchi, 2007), indicating that an impaired immune response in the diabetic context induced by PHA is affected by more than IL-2 signalling. Little is known about the precise signalling pathways and intracellular processes stimulated by PHA and ConA in humans and pigs. However, the impaired immune response selectively induced by PHA in hyperglycaemic conditions, which is not addressed by polyclonal T cell mitogen ConA (Rodríguez-Gómez et al., 2019), points to PHA-associated cell signalling pathways specifically altered in the diabetic condition. In murine T cells, ConA was shown to bind to surface glycoproteins and with high affinity to the co-stimulatory molecule CD28, leading to enhanced T cell proliferation (Ando et al., 2014). If mechanisms hold true for porcine T cells, hyperglycaemia does not affect CD28 signalling in diabetic pigs. In contrast, PHA is able to mimic the stimulation of T cells via crosslinking of T cell receptor and CD3 complex, which is followed by a sustained calcium influx as an important signal for T cell proliferation (Lin et al., 2016). In the Jurkat T cell line, high glucose levels were shown to inhibit T cell activation via an increase in non-enzymatic glycation of concentration-regulating calcium channels and delayed specific activation of these cells with anti-CD3 monoclonal antibody (Boldizsár, Berki, Miseta, & Németh, 2002). Since alterations of calcium homeostasis were shown in various cell types of diabetic patients (Klec, Ziomek, Pichler, Malli, & Graier, 2019), insufficient calcium-mediated signalling in T cells of diabetic pigs might point to impaired PHA-induced proliferation. Thus, the immune cell calcium homeostasis of diabetic *INS*^C94Y^ tg pigs merits further investigations in future studies to examine possible correlations to the impaired T cell response in the diabetic cases observed in our study.

To gain deeper insights into hyperglycaemia-associated molecular changes of CD4^+^ T cells, we analysed the proteome of these cells from diabetic pigs and wildtypes as a hypothesis-generating approach for the dysregulated immune responses. Notably, we obtained a comprehensive proteomic data set for porcine CD4^+^ T cells with a high resolution of 2,704 identified proteins. Among these, we found a total of 3% (80 proteins) with significantly different protein abundances between both groups, indicating distinct proteome differences in CD4^+^ T cells from early stage diabetic pigs, functionally associated to DNA damage responses and cell protection, actin cytoskeleton organization and regulation of cellular energy metabolism. While many studies of immune cells from diabetic type 2 patients were performed by a 2D-gelelectrophoresis approach, only significantly altered spots were analysed subsequently with mass spectrometry (Giorgi et al., 2017; Soongsathitanon et al., 2019). To our knowledge, there is no in-depth proteomic fingerprint of CD4^+^ T cells in hyperglycaemic conditions at initial stage of non-immune-mediated cases of diabetes mellitus. Interestingly, the protein with the highest statistical significance and a 2-fold higher abundance in diabetic pigs was ANXA1. This protein identification corresponds to findings in type 1 (Purvis et al., 2018) and type 2 (G. S. D. Purvis, M. Collino, et al., 2019) diabetic patients, where ANXA1 was previously found with increased levels in serum and plasma, pointing to a pivotal, disease-associated alteration of this protein. However, the biological consequence of elevated ANXA1 in diabetes mellitus remains unexplained (Purvis, Solito, & Thiemermann, 2019). Early studies in mice have shown that ANXA1 acts as a molecular T cell tuner by increasing the strength of T cell receptor signaling and T cell activation (D’Acquisto et al., 2007). Overexpressed ANXA1 induced the differentiation of naïve T cells to pro-inflammatory Th1 cells with a high expression of INFγ (Huang, Zhou, Liu, & Zhang, 2016), promoting an inflammatory milieu. Thus, the higher abundance of ANXA1 in CD4^+^ T cells of diabetic pigs might indicate an enhanced T cell activation with functional importance of a pro-inflammatory immune response. However, also the opposite effect was shown in ANXA1-deficient mouse CD4^+^ T cells (Yang et al., 2013). These cells increased their activation, proliferation and inflammation in the absence of ANXA1, suggesting that this protein rather has an anti-inflammatory role and attenuates an exacerbated inflammatory response (Yang et al., 2013). One possible reason is that externalized ANXA1 regulates the extracellular regulated protein kinases (ERK)/mitogen-activated protein kinases (MAPK) signaling pathway via binding to its formyl peptide receptor (FPR) (Purvis et al., 2018). This pathway is important for T cell activity and strongly contributes to T cell activation and proliferation (Liu et al., 2019) and was shown to be activated in hyperglycaemia (Purvis et al., 2018). Treatment of *Anxa1*^−/−^ mice with human recombinant ANXA1 attenuated inflammatory processes in tissues by reducing MAPK signaling (Purvis et al., 2018). Thus, a higher abundance of ANXA1 in diabetic CD4^+^ T cells may temper signaling to protect the cell from hyperresponsiveness initiated by chronic hyperglycaemia and, thus, exert anti-inflammatory effects. Although we cannot yet define the precise role of ANXA1 in CD4^+^ T cells in the porcine diabetic model, the identification of ANXA1 in humans and pigs, independent of aetiology, is an important finding which underscores a crucial role of this protein in T cell impairment accompanying diabetes mellitus. It furthermore highlights the potential translational quality of this animal model for diabetes research, which is, in our opinion, recommended for future studies to examine the role of this molecule for the observed dysregulated immune responses. Interestingly, one of the most over-represented pathways of altered proteins in CD4^+^ T cells of diabetic pigs was lipophagy. Lipophagy is linked to metabolic dysfunctions of cells (Kounakis et al., 2019; Tian et al., 2019), and thus, pointed to an altered metabolic phenotype in diabetic pigs. We therefore characterised the metabolic immune cell phenotype of PBMC, since cellular metabolism can be a primary driver and regulator of immune cell function (Olenchock et al., 2017). Metabolic properties of diabetic PBMC in this study revealed an increase in mitochondrial respiration and basal glycolytic activity. We found significant increases in mitochondrial oxygen consumption rate (OCR) in PBMC of diabetic pigs. Interestingly, in line with our findings, increased oxygen consumption was also demonstrated in PBMC of type 2 diabetes patients (Hartman et al., 2014). Previously, this group also measured a higher production of reactive oxygen species (ROS) in these cells and hypothesised mitochondrial damage in diabetic PBMC caused by enhanced oxidative stress (Widlansky et al., 2010). Since ROS synthesis is essential for physiological immune activation of lymphocytes but lead to cellular damage if concentration surpasses (Alfatni et al., 2020), this parameter should be determined in PBMC of diabetic pigs, in order to evaluate whether immune cells are impaired by oxidative cell stress. On the contrary, mitochondrial respiration with increased oxygen consumption is used by immune cells to supplement increased glycolysis, which is the main metabolic pathway fueling effector function upon T cell activation (Gaber, Chen, Krauss, & Buttgereit, 2019). Besides increased oxygen consumption rates, we also found a higher basal glycolytic activity in early diabetic pigs, suggesting that hyperglycaemia induced metabolic adaption to local conditions of immune cells and promoted a more activated status in T cells (Bonacina, Baragetti, Catapano, & Norata, 2019). A rapid increase in glycolysis is an important determinant in cellular energy production and required for effective cell proliferation, size expansion, and differentiation (Gaber et al., 2019). Hence, it is also possible that PBMC in young diabetic pigs are highly metabolically active to challenge hyperglycaemic conditions. Our findings indicate that metabolic disturbances occurring at early stage diabetes mellitus, as observed in *INS*^C94Y^ tg pigs, likely cause metabolic reprogramming in PBMC and may also influence their activation status. Since further studies are absolutely required for a better understanding of bioenergetic profiles of PBMC in diabetes mellitus, diabetic pigs enable to define mitochondrial properties and cellular energy production in these cells at initial stage of disease.

Taken together, our findings point to an early, fundamental dysregulation of immune cells when exposed to prolonged high blood glucose. Chronic hyperglycaemia leads to profound molecular and functional changes of T cells reasoned by proteomic and metabolic alterations, and, moreover, affects T cell immune responses *in vitro.* Diabetic pigs qualify for additional translational experiments to explore the crucial role of altered ANXA1 levels in adaptive immunity in diabetes mellitus. The altered metabolic properties of immune cells in young diabetic pigs provided novel information on distinct immune cell metabolism at initial stage of disease. The exact meaning of this significantly altered metabolic phenotype of the immune cells is not known so far, thus, *INS*^C94Y^ tg pigs are a valuable source to further analyse respective changes for the diagnosis, or treatment or even prevention of early stage diabetes mellitus and its immune system-related complications.

## Acknowledgements

The authors thank F. Stetter, M. Weigand, M. Schilloks and B. Amann from Chair of Physiology, LMU Munich for critical discussions and S. Zettler and A. Hinrichs from Chair for Molecular Animal Breeding and Biotechnology, LMU Munich for assistance in blood withdrawal.

This manuscript previously appeared as a preprint on bioRxiv.org (see References).

## Contribution statement

CD conceived and designed the experiments; IMG, SH and CD performed the experiments; IMG and SH analysed the data; SR and EW developed and characterized the *INS*^C94Y^ tg pig model and contributed reagents and materials; IMG and CD wrote the manuscript. All coauthors critically read the manuscript and approved the final version to be published. CD is the guarantor of this work and, as such, had full access to all the data in the study and takes responsibility for the integrity of the data and the accuracy of the data analysis.

## Data availability

The data are available on request from the authors.

