## Supplementary Table 1 for "Chronic hyperglycaemia drives functional impairment of lymphocytes in diabetic *INS*^C94Y^ transgenic pigs"

**Supplementary Table 1. Eighty proteins revealing significant differences in protein abundance in $CD4^+$ T cells of $INS^{C94Y}$ transgenic pigs and non-transgenic wild-type littermates (quantified with at least two unique peptides).** Column 1 (Protein ID) refer to the accession number of the identified protein and column 2 (Description) to the respective protein name, both as listed in the Ensembl protein database (http://www.ensembl.org). Column 3 (Gene name) contains the name of the human orthologue gene and column 4 (Ratio) the ratio of protein abundance between $INS^{C94Y}$ transgenic pigs and wild-type littermates. Column 5 (p-value) shows the p-value, calculated by Student's *t*-test and column 6 contains the number of unique peptides used for quantification.

| **Protein ID** | **Description** | **Gene name** | **Ratio** | **p-value** | **Peptide counts** |
|---|---|---|---|---|---|
| ENSSSCP00000005658 | annexin A1 | ANXA1 | 2,0 | 0,00001 | 2 |
| ENSSSCP00000020292 | WD repeat domain 5 | WDR5 | 0,9 | 0,00001 | 10 |
| ENSSSCP00000018967 | profilin 1 | PFN1 | 1,2 | 0,0005 | 11 |
| ENSSSCP00000009979 | replication factor C (activator 1) 3, 38kDa | RFC3 | 0,8 | 0,0012 | 3 |
| ENSSSCP00000010743 | SWI/SNF related, matrix associated, actin dependent regulator of chromatin, subfamily b, member 1 | SMARCB1 | 0,9 | 0,0021 | 9 |
| ENSSSCP00000027087 | Splicing factor 3B subunit 3 | SF3B3 | 0,9 | 0,0035 | 5 |
| ENSSSCP00000003489 | polymerase (DNA directed), delta 1, catalytic subunit | POLD1 | 0,8 | 0,0039 | 10 |
| ENSSSCP00000012391 | DnaJ (Hsp40) homolog, subfamily C, member 13 | DNAJC13 | 0,7 | 0,0040 | 11 |
| ENSSSCP00000006200 | small nuclear ribonucleoprotein polypeptide A' | SNRPA1 | 0,9 | 0,0041 | 13 |
| ENSSSCP00000003948 | splicing factor 3a, subunit 3, 60kDa | SF3A3 | 0,9 | 0,0044 | 22 |
| ENSSSCP00000011514 | Parafibromin | CDC73 | 0,8 | 0,0053 | 6 |
| ENSSSCP00000024338 | Splicing factor 3B subunit 3 | SF3B3 | 0,9 | 0,0062 | 10 |
| ENSSSCP00000007747 | RALY heterogeneous nuclear ribonucleoprotein | RALY | 0,8 | 0,0065 | 11 |
| ENSSSCP00000018319 | DEAD (Asp-Glu-Ala-Asp) box helicase 42 | DDX42 | 0,9 | 0,0082 | 24 |
| ENSSSCP00000020257 | SNW domain containing 1 | SNW1 | 0,9 | 0,0083 | 11 |
| ENSSSCP00000018316 | SWI/SNF related, matrix associated, actin dependent regulator of chromatin, subfamily d, member 2 | SMARCD2 | 0,9 | 0,0089 | 8 |
| ENSSSCP00000017581 | protection of telomeres 1 | POT1 | 0,9 | 0,0093 | 4 |
| ENSSSCP00000010038 | Sus scrofa esterase D (ESD), mRNA. | ESD | 1,6 | 0,0102 | 13 |
| ENSSSCP00000003670 | protein DJ-1 | PARK7 | 1,2 | 0,0106 | 11 |
| ENSSSCP00000018966 | enolase 3 (beta, muscle) | ENO3 | 1,1 | 0,0142 | 2 |
| ENSSSCP00000018178 | eukaryotic translation initiation factor 4A3 | EIF4A3 | 0,9 | 0,0157 | 13 |

| | | | | | |
|---|---|---|---|---|---|
| ENSSSCP00000002662 | Putative E3 ubiquitin-protein ligase UBR7 | UBR7 | 1,1 | 0,0171 | 3 |
| ENSSSCP00000005624 | SWI/SNF related, matrix associated, actin dependent regulator of chromatin, subfamily a, member 2 | SMARCA2 | 0,7 | 0,0172 | 8 |
| ENSSSCP00000007572 | 5'-3' exoribonuclease 2 | XRN2 | 0,7 | 0,0174 | 13 |
| ENSSSCP00000015761 | DNA polymerase delta subunit 3 | POLD3 | 0,8 | 0,0183 | 4 |
| ENSSSCP00000010279 | cell cycle and apoptosis regulator 2 | CCAR2 | 0,9 | 0,0188 | 31 |
| ENSSSCP00000026458 | ribosomal RNA processing 9, small subunit (SSU) processome component, homolog (yeast) | RRP9 | 0,8 | 0,0189 | 8 |
| ENSSSCP00000010183 | ssc-mir-7144 | ARGLU1 | 0,9 | 0,0197 | 3 |
| ENSSSCP00000018340 | myosin, light chain 4, alkali; atrial, embryonic | MYL4 | 2,4 | 0,0199 | 8 |
| ENSSSCP00000021692 | nitric oxide synthase interacting protein | NOSIP | 0,7 | 0,0201 | 8 |
| ENSSSCP00000003906 | penta-EF-hand domain containing 1 | PEF1 | 0,9 | 0,0203 | 3 |
| ENSSSCP00000017743 | polymerase (DNA directed), delta 2, accessory subunit | POLD2 | 0,8 | 0,0209 | 7 |
| ENSSSCP00000020527 | protein tyrosine phosphatase, non-receptor type 11 | PTPN11 | 1,1 | 0,0210 | 21 |
| ENSSSCP00000023675 | replication factor C (activator 1) 2, 40kDa | RFC2 | 0,8 | 0,0215 | 4 |
| ENSSSCP00000023668 | WD_REPEATS_REGION domain-containing protein | EIPR1 | 1,4 | 0,0217 | 3 |
| ENSSSCP00000024485 | Sus scrofa heterogeneous nuclear ribonucleoprotein C (C1/C2) (HNRNPC), transcript variant 1, mRNA. | EIF4H | 0,9 | 0,0226 | 14 |
| ENSSSCP00000002940 | splicing factor 3b, subunit 3, 130kDa | SF3B3 | 0,9 | 0,0235 | 17 |
| ENSSSCP00000010752 | protein phosphatase, Mg2+/Mn2+ dependent, 1F | PPM1F | 1,2 | 0,0241 | 7 |
| ENSSSCP00000000036 | Tubulin-tyrosine ligase-like protein 12 | TTLL12 | 1,1 | 0,0248 | 27 |
| ENSSSCP00000000601 | cytidine monophosphate N-acetylneuraminic acid synthetase | CMAS | 0,9 | 0,0252 | 15 |
| ENSSSCP00000024473 | enolase 1, (alpha) | ENO1 | 1,1 | 0,0253 | 12 |
| ENSSSCP00000014818 | GATA zinc finger domain containing 2A | GATAD2A | 0,9 | 0,0266 | 9 |
| ENSSSCP00000016213 | retinoblastoma binding protein 5 | RBBP5 | 0,9 | 0,0274 | 9 |
| ENSSSCP00000004229 | peroxiredoxin 1 | PRDX1 | 1,2 | 0,0290 | 9 |
| ENSSSCP00000027721 | protein kinase C, beta | PRKCB | 1,1 | 0,0293 | 4 |
| ENSSSCP00000021929 | fission 1 (mitochondrial outer membrane) homolog (S. cerevisiae) | FIS1 | 0,9 | 0,0296 | 6 |
| ENSSSCP00000000038 | protein kinase C and casein kinase substrate in neurons 2 | PACSIN2 | 1,1 | 0,0312 | 15 |
| ENSSSCP00000021684 | perilipin 3 | PLIN3 | 1,3 | 0,0313 | 7 |
| ENSSSCP00000016029 | hypoxia up-regulated 1 | HYOU1 | 0,9 | 0,0317 | 33 |
| ENSSSCP00000003296 | 40S ribosomal protein S19 | RPS19 | 0,9 | 0,0320 | 7 |
| ENSSSCP00000000849 | Sus scrofa interleukin-1 receptor-associated kinase 4 (IRAK4), mRNA. | IRAK4 | 1,2 | 0,0322 | 2 |
| ENSSSCP00000003309 | X-ray repair complementing defective repair in Chinese hamster cells 1 | XRCC1 | 0,8 | 0,0347 | 5 |
| ENSSSCP00000009333 | DEAH (Asp-Glu-Ala-His) box helicase 15 | DHX15 | 0,9 | 0,0348 | 39 |
| ENSSSCP00000010920 | cell division cycle and apoptosis regulator 1 | CCAR1 | 0,9 | 0,0357 | 8 |

| | | | | | |
|---|---|---|---|---|---|
| ENSSSCP00000000041 | polymerase (DNA-directed), delta interacting protein 3 | POLDIP3 | 0,8 | 0,0362 | 6 |
| ENSSSCP00000008216 | Sus scrofa heat shock 27kDa protein 1 (Hsp27), mRNA. | HSPB1 | 1,2 | 0,0367 | 10 |
| ENSSSCP00000007783 | RNA-binding protein 39 | RBM39 | 0,9 | 0,0376 | 16 |
| ENSSSCP00000010175 | \|11\|ENSSSCG00000009528\|ENSSSCT00000010446 | TPP2 | 1,1 | 0,0384 | 18 |
| ENSSSCP00000018643 | Sus scrofa non-metastatic cells 2, protein (NM23B) expressed in (NME2), mRNA. | NME2 | 1,1 | 0,0387 | 5 |
| ENSSSCP00000023248 | small nuclear ribonucleoprotein D2 polypeptide 16.5kDa | SNRPD2 | 0,9 | 0,0400 | 5 |
| ENSSSCP00000003447 | Sus scrofa interferon regulatory factor 3 (IRF3), mRNA. | IRF3 | 1,2 | 0,0417 | 7 |
| ENSSSCP00000006744 | Sus scrofa microsomal glutathione S-transferase 3 (MGST3), mRNA. | MGST3 | 0,7 | 0,0425 | 5 |
| ENSSSCP00000005485 | mitochondrial ribosomal protein S11 | MRPS11 | 1,4 | 0,0429 | 3 |
| ENSSSCP00000025894 | NCK adaptor protein 1 | NCK1 | 1,1 | 0,0437 | 5 |
| ENSSSCP00000008042 | protein phosphatase 1, regulatory subunit 3D | PPP1R3D | 1,6 | 0,0440 | 4 |
| ENSSSCP00000017561 | ATPase, H+ transporting, lysosomal 14kDa, V1 subunit F | ATP6V1F | 1,1 | 0,0441 | 2 |
| ENSSSCP00000025052 | carbamoyl-phosphate synthetase 2, aspartate transcarbamylase, and dihydroorotase | CAD | 1,1 | 0,0453 | 15 |
| ENSSSCP00000018067 | cytoplasmic FMR1 interacting protein 2 | CYFIP2 | 0,8 | 0,0455 | 18 |
| ENSSSCP00000018929 | chromosome 17 open reading frame 85 | NCBP3 | 0,8 | 0,0461 | 2 |
| ENSSSCP00000003276 | Platelet-activating factor acetylhydrolase IB subunit alpha | PAFAH1B 3 | 1,1 | 0,0469 | 9 |
| ENSSSCP00000023178 | interferon-induced protein 35 | IFI35 | 0,9 | 0,0479 | 3 |
| ENSSSCP00000027336 | mutL homolog 1 | MLH1 | 0,8 | 0,0484 | 6 |
| ENSSSCP00000013941 | Sus scrofa PRP19/PSO4 pre-mRNA processing factor 19 homolog (S. cerevisiae) (PRPF19), mRNA. | PRPF19 | 0,9 | 0,0491 | 19 |
| ENSSSCP00000003451 | SR-related CTD-associated factor 1 | SCAF1 | 1,3 | 0,0506 | 3 |
| ENSSSCP00000018184 | chromobox homolog 8 | CBX8 | 0,8 | 0,0509 | 2 |
| ENSSSCP00000028171 | poly-U binding splicing factor 60KDa | PUF60 | 0,9 | 0,0512 | 13 |
| ENSSSCP00000010503 | Sus scrofa protein kinase, AMP-activated, beta 1 non-catalytic subunit (PRKAB1), mRNA. | PRKAB1 | 1,3 | 0,0517 | 2 |
| ENSSSCP00000010155 | importin 5 | IPO5 | 1,1 | 0,0524 | 35 |
| ENSSSCP00000025726 | Rho GTPase activating protein 17 | ARHGAP1 7 | 1,2 | 0,0544 | 5 |
| ENSSSCP00000011466 | O-6-methylguanine-DNA methyltransferase | MGMT | 0,9 | 0,0545 | 4 |
