## Supplementary Figure 1 for "Chronic hyperglycaemia drives functional impairment of lymphocytes in diabetic *INS*^C94Y^ transgenic pigs"

**Supplementary Figure 1. Lymphocyte subpopulations in porcine PBMC.** Scatter plots illustrate percentages of lymphocyte subpopulations in wild-types and $INS^{C94Y}$ tg pigs. Flow cytometry analyses revealed no significant differences in the percentage of $CD3^{+}$ T cells (wt n=19; tg n=16) **(A)** and $CD79a^{+}$ B cells (wt n=7; tg n=7) **(B)** and furthermore no significant differences in T cell subsets $CD4^{+}$ (wt n=19; tg n=17) **(C)**, $CD8\alpha^{+}$ (wt n=20; tg n=17) **(D)**, $CD4^{+}CD8\alpha^{+}$ (wt n=7; tg n=8) **(E)** and $SWC5^{+}$ γδ T cells (wt n=10; tg n=11) **(F)**.

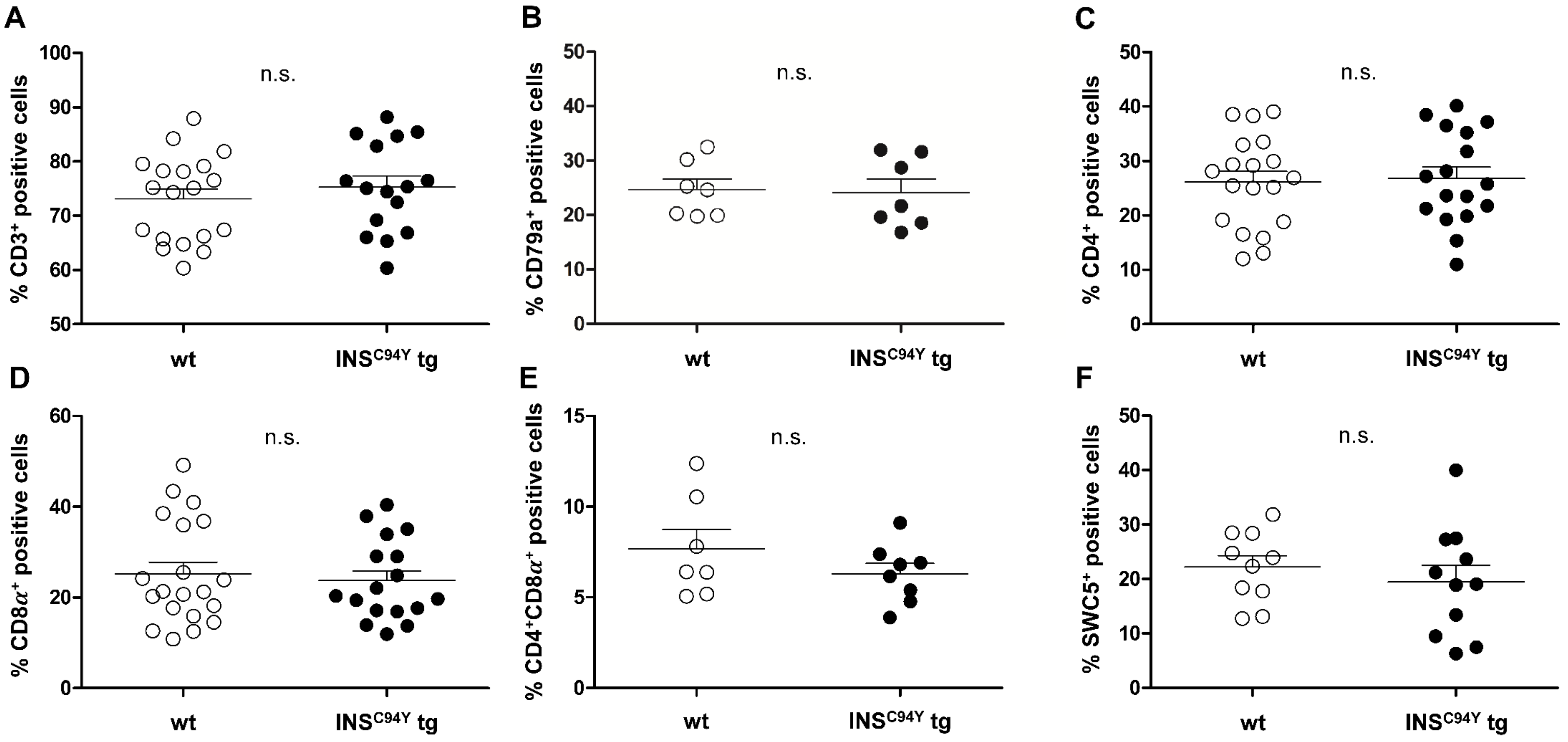
